# The temporal dynamics of the Stroop effect from childhood to young and older adulthood

**DOI:** 10.1101/2021.08.13.456209

**Authors:** E. Ménétré, M. Laganaro

## Abstract

It is well admitted that children and older adults tend to show longer response latencies at the Stroop task than young adults. The present study aims at clarifying the rational of such changes from childhood to adulthood and in ageing by comparing the impacted cognitive processes across age groups. More precisely, the aim was to clarify if all processes take more time to be executed, hence implying that longer latencies rely mainly on processing speed or if an additional process lengthens the resolution of the conflict in children and/or older adults. To this aim we recorded brain electrical activity using EEG in school-age children, young and older adults while they performed a classic verbal Stroop task. To decompose the signal in the underlying brain networks, we used microstates analyses and compared congruent, incongruent and neutral trials across the three age-groups. Behaviorally, children and older adults presented longer latencies and larger Stroop effects relative to young adults. The microstates results showed that children tend to present different brain configurations compared to both adult groups, even though some brain configurations remained identical among the three groups. In particular, additional brain networks were involved in children to perform the Stroop task, which party account for the longer latencies in this group. By contrast, in aging the results favor the general slowing hypothesis rather than a decline in a specific process since all involved brain networks were similar in the two adult groups but slowed down in the older one.

## 1. Introduction

Attentional control is probably one of the most studied topics in cognitive psychology. Historically, this process has been investigated with many paradigms among which the Stroop task [1], extensively used. Even though the exact design of the task varies across studies, the Stroop paradigm can generally be defined as the presentation of two congruent or incongruent pieces of semantically related information while the subject has to “name” (manually or orally) only one of them, usually the color of the ink in which words are displayed. It is typically observed that subjects are slower to process incongruent items (e.g. naming the ink color of the word “blue” written in red) than congruent ones (e.g. naming the ink color of the word “green” written in green), and the congruent condition seems to lead to less errors than the incongruent one [2]. This difference in reaction times and accuracy is known as the *Stroop effect*. Since 1935, a large number of studies have tried to understand the core processes underlying the Stroop effect, namely how the conflictual information of the word reading and color naming processes is handled from a cognitive and neural point of view. Authors globally agree on a two levels of conflict model explaining the Stroop interference. The first one appears after the perception of an incongruency in the stimulus (stimulus conflict), while the second one is a response-based conflict, occurring when the response has to be selected among the two possible answers [3-6]. Moreover, the Stroop task involves some additional processes relative to other attentional control tasks such as the flanker or the Simon tasks. Indeed, it involves components engaged in word production as well as executive processes allowing to select the correct answer [7].

To go beyond the conclusions obtained from behavioral studies, researchers used neuroimaging approaches, such as magnetoencephalography - MEG - [8,9] and electroencephalography - EEG - [10-21], allowing to infer on the cognitive processes and their dynamics. The results of these electro- and magnetophysiological studies converge on some central conclusions. Indeed, most studies reported a more negative deflection for incongruent relative to congruent trials around 400 to 450ms associated with negative centroparietal topographies (referred to as N400 hereafter). A second component has also been described for the same comparison (congruent versus incongruent trials) at a later time-period. The latter sometimes is referred to as the slow potential component (SP), late positive complex (LPC), the P600 or SP600 (SP600 hereafter), and is widely spread around 600ms, showing a centroparietal positivity. Notwithstanding converging results on the components underlying the Stroop effect, the cognitive processes behind these components have been widely debated. A general agreement has however emerged on the idea that the N400 is associated to the detection of the conflict (conflict at the stimulus level) while the second component has been associated with conflict resolution (conflict at the response level).

It is also noteworthy to mention that the vast majority of behavioral and neuroimaging studies used manual responses (mapping between the color response and a keyboard key), and very few studies had participants giving a verbal response [8,18,21-23]. These two response modalities do not always lead to the same results. Some studies tried to understand the electrophysiological differences characterizing the verbal and manual versions of the task. Initially, Liotti and colleagues (2000) confirmed the behavioral finding already reported in earlier studies that the Stroop effect was larger in the verbal modality [2] and showed increased N400 and SP600 effects. However, when comparing the response latencies with the timing of the components, it appears that the SP600 effect is located at the same temporality as the response time. It implies that the observed difference on the SP600 component could simply be attributable to a shift in the response artifact latencies between the congruent and incongruent conditions. Classically, to overcome this limitation, the ERP signal is aligned first to the stimulus (stimulus-aligned signal) and then to the response (starting the temporal window from the onset of the response and extracting anterior time points, i.e. response-aligned signal) [24,25]. By using this procedure, some authors showed significant differences between congruent and incongruent conditions only for the vocal version of the task close to the production of the response [23]. These results were interpreted as an interference effect appearing not only at a semantical or lexical level, but also at the word form encoding level. In other words, the verbal and manual Stroop tasks are comparable in terms of involved cognitive processes except that the verbal Stroop task involves also a word form encoding conflict component explaining increased interference effect for this modality.

To sum up, the ERP components characterizing the cognitive processes involved in the Stroop task are globally well defined and their underlying mental processes are consistently described. However, this is only the case for young adults: other populations such as typical children and healthy older adults might show differences in mental processes and brain activations when performing the Stroop task. As further detailed below, the literature shows that the timing of the ERP components elicited by the task is modulated by the age of the participant. It is nonetheless still unclear (1) how the Stroop effect evolves across the lifespan both behaviorally and neurophysiologically and (2) if the involved processes are identical at all ages. Hereafter we will review the available evidence on these two issues and the questions that are still debated or unresolved.

### 1.1 The Stroop effect across the lifespan

The large majority of what we know about the Stroop task comes from studies based on young adult participants. A few studies have investigated changes in executive control over the entire lifespan with the Stroop task or some other tasks [17,26–30], but these studies are not numerous in the wide literature of the evolution of executive control abilities. That is why in the following, we will first examine the evolution of the Stroop effect from childhood to adulthood, and at a second stage, focus on the mechanisms explaining the decrease of performances in aging.

#### 1.1.1. From childhood to adulthood

The literature on the evolution of performance from childhood to adulthood suggests that even after a short exposure to reading, children show a strong and reliable Stroop effect despite longer latencies and an increased error rate [2,31]. In school-age children, the Stroop effect evolves in a non-linear pattern, which seems entirely related to reading acquisition as when accounting for reading level, a linear trend appears [32]. This observation involves that the development of attentional control and reading abilities interact in the evolution of the performances on the Stroop task across childhood. Using an ERP approach with children aged from 6 to 12 years old performing a semantic Stroop task (e.g. naming the font of the word “sky” written in blue), Jongen and Jonkman (2008) claimed that stimulus conflict (reflected by the N400 component) appears early in the development, which translates by an early setup of the inhibition mechanisms, while the response conflict (reflected by the SP600) continues to mature until 12 years old. This result is in accordance with a literature review, stressing that the processes at play when handling interference are not fully developed before the age of 12 years or later [34], as well as a study investigating the evolution of the conflict adaptation (adaptation of the attentional control system from one trial to the next) [35]. The literature on other attentional control tasks such as the flanker task suggests that conflict detection processes reach their maturation at the age of 10 years old while, surprisingly, conflict resolution does not seem to evolve. The absence of evolution of conflict resolution processes could be related to the non-verbal character of the task [36]. Overall, authors conclude that since the ERP components are already present in childhood, processes are equivalent but differ in their maturational status. However, even though processing speed evolves during development, it cannot explain the entire changes observed between childhood and adulthood [37].

#### 1.1.2 Aging

At the other end of the lifespan, it is well admitted that cognitive performances decline, and this is particularly well described regarding conflict processing abilities. Different explanations have been proposed regarding the general alteration of cognitive performances with aging. Two hypotheses have been suggested: the general slowing hypothesis and a specific alteration of executive functions with aging. These hypotheses were derived partly from the decline observed in the Stroop task.

The general slowing hypothesis explains the longer latencies in aging by a slowdown of all processes. The decline could then simply be explained by a decrease in attentional resources but a stability of the inhibition abilities [38-42]. Applied to the Stroop task, this would imply that both stimulus and response conflicts are impacted in the same way by aging, since all implicated cognitive processes are slowed down to compensate the needed increased effort. Mathematical models of the effect of aging on the Stroop interference concluded that the general slowing hypothesis explained all the effect, leaving no room for the deterioration of a specific process [41,42]. Likewise, more recently, a meta-analysis on several inhibition tasks, concluded that only the general slowing hypothesis explains the observed changes [39]. On top of these mathematical and statistical considerations regarding the Stroop effect evolution, an experimental approach manipulating the inter-stimuli delay showed that by allowing participants more time to rest, elderly subjects reach the performance of their young counterparts [38].

On the other hand, the second hypothesis claims that the main explanation for the longer latencies and increased interference effect in elderly compared to young adults is related to the decline of (a) specific cognitive process/es in aging that cannot be explained by general slowing [26,43-49]. Regarding the Stroop task, this hypothesis might imply a different alteration of one or several process(es) (conflict detection or conflict resolution) more than others. This effect was shown by combining approaches of semantic Stroop task and single-colored-letter paradigms (i.e. only the central letter of the color word is displayed in color). Some authors reported a deficit in aging only in the response conflict but not in the stimulus conflict (or semantic conflict) [49]. From a neural point of view, a princeps ERP study using the Stroop task in old adults showed a modulation of the P300 component (positive component peaking around 300ms after stimulus onset and showing a centro-parietal positivity), as well as the N400, SP600 and error-related negativity components [12]. The results were interpreted as a specific decline of these mechanisms in aging, although the authors did not conclude on the origin of such decline. It seems that both stimulus and response conflict are impacted since the N400 as well as the SP600 are modulated, nevertheless further clarifications are needed to better characterize this phenomenon and to disentangle the general slowing from specific deficit theories.

At a first glance the two theories are contradictory, but it is possible that they are two faces of the same coin. All processes can be slowed down in aging due to attentional decline but some may be affected more than others, reflecting a decline in specific executive processes. Moreover, a slowdown in latencies does not exclude that compensatory mechanisms are involved. The complementarity of the two theories has been investigated using behavioral approaches [50]. Applying a hierarchical regression model, the authors observed that processing speed (measured by a simple reaction time task) accounted for a large part of the variance but age explained a specific part of the residual variance, supposing a specific deficit in executive functions, although this result may be further clarified using electrophysiological approaches.

To sum up, state of the art literature agrees that brain activations change from childhood to aging. Indeed, components measured on the EEG signal vary across ages in amplitude and timing of the peaks. The changes in latencies and error rates in the Stroop task are generally interpreted as the consequences of less developed processes in childhood compared to adulthood and a decline in these same processes with aging. Differences in timing could translate a linear development of attentional control as well as a linear decline of the performances of the latter, but no arguments were highlighted in the neurophysiological literature regarding a specific decline of attentional control, as argued partly in the behavioral literature.

### 1.2. A single mechanism underlying congruent and incongruent trials, at least in adulthood

One of the core questions tackled in the first neuroimaging literature dedicated to the Stroop paradigm was to find how the conflict is processed and whether it requires an additional mechanism as compared to congruent or neutral trials. EEG results usually indicate similar components across conditions, except for a difference in amplitude of the N400 component and the SP600 (but see concerns about the SP600 in Stroop tasks with manual - button press - responses reported by Zahedi and colleagues (2019) and summarized above). Since latencies vary between the congruent and incongruent conditions, the amplitude difference observed between the components might be attributed to a shift in the signal rather than to different processes (i.e. differences in amplitude) underlying the two conditions. The issue of whether there is a pure shift of the same components or different brain processes across conditions is best investigated using microstates analyses. This method allows to investigate the presence of different global electric fields in the signal, i.e. of different underlying mental processes [51-54], but also their relative onset and duration. The analysis would allow to further describe whether the same processes underpin the two conditions in the Stroop task and whether this is the case across age-groups. Several studies investigated the Stroop effect using microstates [16,55-57], but to our knowledge, only two of them have investigated the brain networks involved in both congruent and incongruent conditions. Khateb and colleagues proposed a modified Stroop task to young adult participants and segmented the ERP signal in periods of quasi-stable global electric fields [16]. Their results confirmed that the brain processes are common between congruent and incongruent trials, but that one microstate was extended in the incongruent trials compared to the other condition, reflecting differences in response latencies. The differences in latencies could thus be explained by the increased duration of a particular topography in the signal of incongruent trials. The Stroop interference would then be recruiting a large network and the results speak against a specific module at play to detect and resolve conflicts, in accordance with the rest of the literature. These results were confirmed by a second study showing the appearance of the same microstates maps for congruent and incongruent trials in the EEG signal, however showing a different distribution of the maps starting at about 500ms until 750ms. The authors performed source localization analyses on this time-window which pointed out significant activation differences located in the ACC [57]. This study hence confirmed what was reported in previous literature, hence that the ACC was sensitive to the time spent on an item rather than the presence of a conflict [7]. According to the conflict monitoring model, this structures activates in presence of conflictual information and acts as a conflict detector [58]. Nevertheless, it has been suggested that the ACC might appear more activated in the incongruent condition given its longer activation time [7,57]. It is worth to mention that the time window carrying the Stroop effect is different in the two mentioned studies investigating the Stroop effect using microstates analyses: Khateb and colleagues showed an elongation of the microstate underlying the N400 while Ruggeri et al. reported a difference in the SP600 time window. The different loci of the effect remain unclear, but might be due to the design of the tasks, since Khateb and colleagues used a passive pseudo-Stroop task.

In the present study, we went a step further by investigating whether the same underlying processes for congruent and incongruent trials are also observed during development and in aging. As summarized above, despite behavioral and ERP changes observed across the lifespan on the Stroop task, the underlying processes seem to be the same, although less efficient during maturation and in aging. This issue has however never been confirmed directly by considering both children and older adults in a same study. To address this, we designed a classical serial Stroop task including congruent, incongruent and neutral items (rows of symbols displayed in different colors) while recording the participants brain activity with high density EEG in school-age children (10 to 12 years old), young adults (20 to 30 years old) and older adults (from 58 to 70 years old). EEG signal was first analyzed with peak and mass univariate tests on the waveform amplitudes, and in a second step, using topographical or microstates analysis. Finally, source localization analysis was performed on the time-window of the microstates carrying the Stroop effect to clarify the underlying brain structure and associated cognitive processes. Even though children, young and older adults were included in the same analysis to distinguish better the different trend of development and aging, the results were interpreted separately, as distinct research questions.

According to the literature on the development of the Stroop effect reported above, we expected that children would have the same brain networks at play when performing a congruent or an incongruent trial. However, it is still unclear if children are slower and display larger Stroop effects relative to young adults because of the immature attentional resources impacting all processes or if only some processes need more time to become fully efficient. In other words, if microstates are identical in children and in young adults but are only different in duration, then it might suggest that the executive processes are already mature in school-age children but attentional resources are not developed enough to allow reaching the performances of young adults. Another possible explanation would be that one specific mechanism is still immature in children and explains the slower reaction times. This hypothesis would be confirmed if additional microstates were identified in children relative to young adults or if a specific microstate was significantly more impacted in children.

The same rationale may be applied regarding aging. A lengthening of all microstates would favor the general slowing hypothesis while a supplementary process (additional microstate) in older adults relative to young adults would favor a compensatory mechanism. Finally, a disproportionately more present process in the older adults group would favor the decline of a specific brain network in aging.

## 2. Method

### 2.1 Participants

Seventy-four participants were recruited, divided into three age groups (children, young adults and older adults). Among them, only participants with a minimum of 25 artifact-free ERP epochs per condition (see pre-analyses) were retained. The final sample thus included 53 participants: 17 children from 10 to 12 years old (mean age: 11; SD = 0.79; 7 females), 18 young adults from 20 to 30 years old (mean age: 23.9; SD = 2.97; 13 females) and 18 older adults from 58 to 70 years old (mean age: 64.9; SD = 3.6; 14 females). This sample size seemed reasonable given that previous studies included from 8 to 25 participants in average [14,18,59]. Older adults aged less than 70 years old were preferred to elderly to avoid cognitive decline and selection bias related to the fact that only the more cognitively performant older persons agreed to participate to neuroscientific studies. All participants were right handed [60] native French-speakers, and did not report any language, neurological, psychiatric or color vision impairment. All participants or their legal representative gave their written consent before the beginning of the procedure and received a financial compensation for their participation. The entire procedure was approved by the local Ethics Committee.

The participants were part of a larger group from which only the behavioral data has been analyzed in previous study [40].

### 2.2 Materials

A 180 trials, classical four colors Stroop task requiring verbal responses was used. The verbal stimuli were four color names in French (“*bleu*”; “*jaune*”; “*rouge*”; “*vert*”, respectively *blue, yellow, red* and *green*) as well as symbols displayed in different colors, were those of Fagot and colleagues (2008). The task set encompassed 60 congruent (the color font and the color words match, e.g. the word “*bleu*” written in blue), 60 incongruent (there is a discrepancy between the color font and the color word, e.g. the word “*vert*” written in red) and 60 neutral items (“++++”; “^^^^”; “’’’’”; “****”) displayed in one of the four different colors. All verbal stimuli were presented in lower case at the center of the screen.

### 2.3 Procedure

Participants sat at approximately 80 cm of a 17 inches computer screen (refreshment rate: 50Hz). Stimuli were presented using the E-Prime software (E-studio). Oral responses were recorded by a microphone and sent to the E-Prime software to be recorded and labelled. To estimate reaction times, the onset of the production was manually retriggered offline using the CheckVocal software [62].

Participants received the instruction to name in which color the stimuli were displayed. They were also asked to produce their response orally as fast and accurately as possible. Regarding the instructions relative to the EEG recording, in order to avoid artifacts in the signal, the participants were asked to remain still and blink only after the end of the production of their responses.

The trial structure was identical for all the age groups. First, a white fixation cross was displayed on a black background for 500ms. To avoid visual responses’ contamination on the temporal window of interest due to the fixation cross, a black screen was presented for 200ms. The stimulus was then presented for 1500ms, followed by a variable interstimuli black screen for which the duration was ranging randomly from 1000 to 1200ms.

Before the beginning of the task, a training phase of 32 items including all the possible stimuli was proposed to the participant. The purpose was first to familiarize the participants with the task, but also to avoid, or at least attenuate, an eventual novelty effect on the first trials.

### 2.4. Behavioral data analysis

Statistical analyses, data wrangling and figures were performed using the R software (V.3.8) [63] with the libraries base, dplyr [64], tidyr [65], ggplot2 [66], lme4 [67], lmerTest [68], emmeans [69], NPL (Ménétré, 2021; available at: https://github.com/EricMenetre/NPL), optimx [71], car [72] and glmmTMB [73] packages. Data and code generating the figures and the analyses is available on the Yareta platform (URL) [AVAILABLE SOON ON THE YARETA PLATEFORM].

A trial was excluded if the participant gave an incorrect color name, corrected himself/herself immediately after the production of an error, or did not give any response. Minor hesitations such as phonological transformation (example: “r-rouge”) were considered as correct. Regarding latencies, all response times under 200ms were excluded, as well as values outranging the threshold of 2.5 standard deviations above and under the mean of the age group. The respective rejected trial rates for congruent, incongruent and neutral condition was: 0.85%, 3.85% and 0.35%.

The data were analyzed using linear mixed models with the lmer function (lme4 package) for latencies and generalized linear mixed models fitting a binomial distribution with the glmer function (from the same package) regarding accuracy. Post-hoc analyses were estimated using the emmeans function performing a Tukey test. It is noteworthy to mention that since the model initially did not converge, the optimization algorithm was changed for “nlminb”. For both models, post-hocs were estimated by a Tukey test targeting relevant comparisons. Moreover, to obtain main effects on generalized linear mixed models, the Anova function from the car package was used instead of the classical anova function from base R used for lmer models.

### 2.5. EEG acquisition and pre-analyses

The electroencephalogram (EEG) was recorded continuously during the task by 128 electrodes placed on a nylon cap following the 10-5 system. The signal was acquired by a Biosemi amplifier including two active electrodes (ActiveTwo system, Biosemi V.O.F. Amsterdam, Netherlands) at a sampling rate of 512Hz including an online DC filter ranging from 0.01 to 104Hz and a 3dB/octave slope.

Regarding data cleaning, all the manipulations were performed with the Cartool software (V. 3.91) [74] and with the R software (V.3.8) [63]. The signal was first filtered by using a notch filter at 50Hz and a zero-phase shift order 2 bidirectional butterworth bandpass filter including a highpass filter at 0.3Hz and a lowpass filter at 30Hz. The signal was then downsampled at 256Hz to reduce the computing time and the number of multiple comparisons of the statistical analyses. A first interpolation was performed on the entire signal for noisy electrodes following a 3-D spline method [75]. The mean number of interpolated electrodes was 4.79 (SD = 1.84) (with at maximum 10 interpolated electrodes, or 7.81% of the 128 electrodes). Regarding the event related potentials (ERP) extraction, two time windows were selected: a stimulus-aligned period and a response-aligned period. The rationale of two alignment points has been presented in the Introduction [24,25]. The data aligned to the stimulus for a shorter time period than the behavioral latencies allow to track the N400 effect; the data aligned to the response onset (backwards) allows the investigation of later stages of word form encoding. Finally, this approach is also adopted to avoid the presence of artifacts due to articulation in the EEG signal. The first period included the first 25 time frames (~100ms) before the stimulus presentation, which was taken as baseline, and the 130 time frames (~520ms) following the stimulus presentation. This stimulus-locked ERPs lasting ~520ms ensured the entire signal to be at least 100ms shorter than the fastest RTs (629ms in the C condition in young adults, see Table 1). The second window was selected backward from the response to the stimulus and lasted 100 time frames (~400ms). For the purpose of the EEG analyses, the response latencies used as onset of the response-aligned period were reduced by 100ms to avoid measuring the premises of articulation [25]. Since no automatic algorithm of artifacts correction were used, all the epochs were visually inspected and only the artifact-free epochs were retained for the averaging, and grouped according to the Stroop conditions (congruent, incongruent and neutral). Finally, a second interpolation was performed on the averaged data of each subject (mean number of interpolated electrodes: 3.38 (SD = 2.75); maximum 9 electrodes or 7.03% of the total number of electrodes) using the same method as described above but avoiding interpolation of previously interpolated electrodes or their direct neighbors. When combining the two interpolations, on average, 8.17 (SD = 3.3) electrodes were interpolated per participant and the maximum of interpolated electrodes was 16, or 12.5% of the total number of electrodes. On average, 17.48% (SD = 10.87), 26.82% (SD = 13.94) and 19.78% (SD = 13.08) of the trials were rejected for the congruent, incongruent, and neutral conditions, respectively, including the ones already rejected during the behavioral analyses.

**Table 1:**
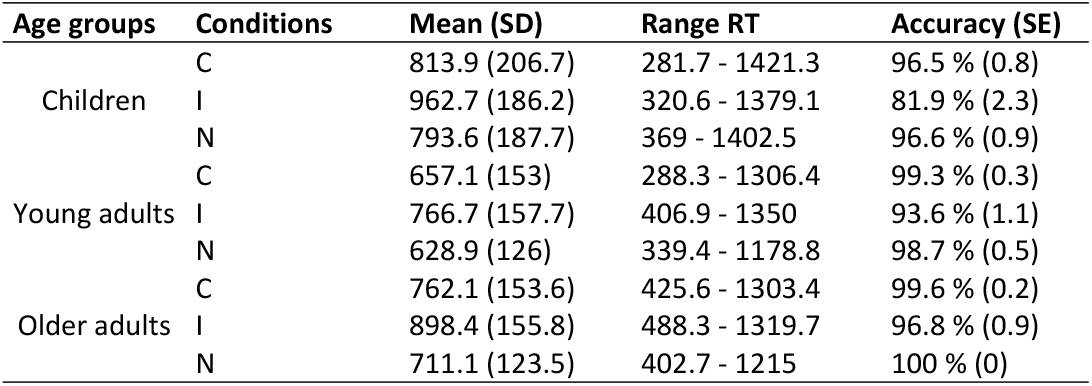
descriptive statistics regarding the mean latencies with their relative standard deviations, the range between the minimum and maximum reaction times observed and accuracy with its standard error.

### 2.6 EEG analyses

To ensure comparison with the rest of the literature, first, the waveforms of the signal was analyzed. This was achieved by a peak analysis and by a mass-univariate statistic. In a second step, the topography of the data was considered. It was verified that the signal was topographically consistent using a topographical consistency test before running an analysis of variance between the three conditions and age groups on the signal (tANOVA). This analysis will guide the interpretation of the microstates and will allow to relate topographical differences with those observed on the waveform analyses. The signal was then segmented in microstates and the characteristics of the microstates (presence, duration and onset) were extracted. Since these microstates tend to be difficult to relate to cognitive processes, source localization analyses were performed to identify the generators behind the most relevant brain networks identified by the microstates analyses.

#### 2.6.1. Waveforms statistics

Mass univariate tests on amplitudes were performed using the STEN software [76]. The data were analyzed using a 3X3 (three conditions, i.e. congruent, incongruent and neutral, by three age groups, i.e. children, young adults and older adults) parametric ANOVA. A spatial and a temporal criteria were applied: only periods of significance of at least 4 time frames (~16ms) on at least 4 electrodes were included. A significance threshold of 0.01 was defined and, since one test per time point was performed, a correction for multiple comparisons was applied. Multiple comparisons was corrected using a Bonferroni approach.

To match and replicate the approaches used in previous studies, a peak analysis was performed. Since literature usually report electrodes from the midline of the EEG cap, we selected the FPz, Fz, Cz, Pz and Oz electrodes. Since response-aligned analyses are rare in the literature, only the stimulus-aligned period was considered for this analysis. The time window retained for the analyses was selected based on visual inspection and ranged from 300 to 420ms. A repeated measures 3(conditions: congruent; incongruent; neutral)X3(groups: children; young adults; older adults) ANOVA was calculated on mean amplitude per subject and condition in the time window. Post hoc decomposition of the interaction was performed using a Tukey test.

#### 2.6.2 Topographic analysis of variance (tANOVA)

Topographic analyses were performed on both (stimulus-aligned and response-aligned) ERP signals across groups using the Ragu software [53,54]. A test evaluating the consistency of the topographies of the averaged epochs was also performed, prior to the interpretation of the tANOVA (topographic ANOVA) results. This test aims at assessing if the signal shows consistent topographies, guarantying the reliability of the topographical analyses such as the tANOVA and the microstates analyses. First, a tANOVA was estimated including the three conditions (3X3 design: three conditions, i.e. congruent, incongruent and neutral, by three age groups, i.e. children, young adults and older adults) and then to investigate the Stroop effect, the contrast between congruent and incongruent conditions was calculated in a separate analysis (2X3: 2 conditions by three age groups).

#### 2.6.3. Microstates analysis

The microstates analysis allows to decompose the EEG/ERP signal in periods of topographical stability (or quasi-stability). It has been proposed that when a cognitive process is taking place, the topography of the scalp EEG remains stable, reflecting the activation of a specific brain network. During this stability, different brain areas are connected according to one or several firing frequency/ies [77-80]. Each of these periods, or microstates, can be segmented from the evoked potentials and are known to reflect the activation of a network, generally underlying one or several cognitive processes.

Microstates analysis was performed separately on stimulus aligned and response aligned ERP signals across groups using the Ragu software [53,54]. The software performed a segmentation on the averaged ERP of each subject using a cross-validation method based on the topographical atomize and agglomerate hierarchical clustering (TAAHC) algorithm, for a user pre-defined user range of topographies (here from one to 30). At each fold (data are segmented in subsamples named folds and models are trained on a majority of the folds and tested on the rest), the segmentation is operated on the training data and the global explained variance is reported on both training and test data. The user then choses the more parsimonious number of template maps explaining the most variance in the signal, usually on the test data to avoid overfitting. For display, the selected maps, after applying a smoothing method (aiming at avoiding small microstates appearing in the middle of another larger one), are fitted on the grand average for each condition and age group. For a more detailed explanation of the procedure, see: [53,81]. It is noteworthy to mention that to avoid fitting microstates related to the P1 later in the signal, the segmentation on the stimulus-aligned data was performed only after the P1 window (73 time frames, or ~192ms). This timing was based on the peak of dissimilarity observed in the children’s grand average (since children presented the wider P1).

The procedure allows to extract quantifiers of the microstates such as presence/absence or duration in terms of number of time frames, onset, offset, center of gravity, etc. per subject. Even though the original paper [53] describing the Ragu software did not include the possibility to extract quantifiers per subjects, microstate and condition, this option has been added recently. Here, only template map presence/absence, onset time and duration were considered for statistical analyses. Regarding the presence/absence, the data were analyzed by generalized linear mixed models (one for the stimulus and another for the response-aligned signal) fitting a binomial distribution. Onset was analyzed by a generalized linear mixed model fitting a gamma distribution using a log link function. Since the duration data followed a Poisson distribution (count of the number of time frames occupied by a microstate in the signal), with an over-representation of the zero value, translating the presence or absence of the topography in the subject, the use of a generalized linear mixed model fitting a zero-truncated Poisson distribution (Hurdle models) was justified. The analyses were performed using the glmmTMB function, and the post-hocs using Tukey tests from the emmeans package. For models which failed to converge (all of them except for the Hurdle model on microstates duration, the optimizer function has been adapted using the optimx package, relying on the “bobyqa” algorithm.

#### 2.6.4 Source localization analysis

Source localization analysis was performed using the Cartool software. MRI from the MNI-152 symmetric atlas for adults (applied to both adults groups) and a pediatric atlas (sample aged from 10 to 14 years) [82] for the children group were used. The entire brain was extracted from the MRI (combining gray and white matter) and electrodes position was coregistered on the MRI head of adults and children templates. The solution space was estimated using 6000 solution points and the lead field was calculated using a LSMAC 4-shell isotropic method. To be noted that the lead field was calculated once per age group since the calculation method takes into account the average age of the group to estimate skull thickness. To select the best inverse solution method, an estimation of the sources underlying the P100 was performed with several inverse models (minimum norm, weighted minimum norm, sDale, sLORETA, LORETA, LAURA, eLORETA), and the one providing the closest activation around the occipital areas was selected. LORETA [83], gave the closest estimation, displaying a bilateral occipital activation without engagement of the cerebellum. Inverse solutions were estimated using an inverse matrix per age group based on the grand average of each conditions in the stimulus- and response-aligned signal and before computing, a spatial filter was applied. The estimation of the sources was performed based on the microstates analysis. For each of the maps of interest, the onset and offset of the map per condition was selected and the corresponding sources from the grand averages per conditions and age groups were calculated. Only the maps highlighting a significant difference in terms of presence or duration among conditions or age groups explaining the Stroop effect were retained.

## 3. Results

### 3.1 Behavioral results

As shown in Table 1, children presented the slowest latencies in all conditions (mean = 851ms; SD = 207), and young adults (mean = 682ms; SD = 157) were faster than the older adults’ group (mean = 788ms; SD = 164). Children also displayed the highest standard deviations regarding errors and reaction times, while young and older adults showed comparable variances despite differences in reaction times, as highlighted by Figure 1.

**Figure 1:**
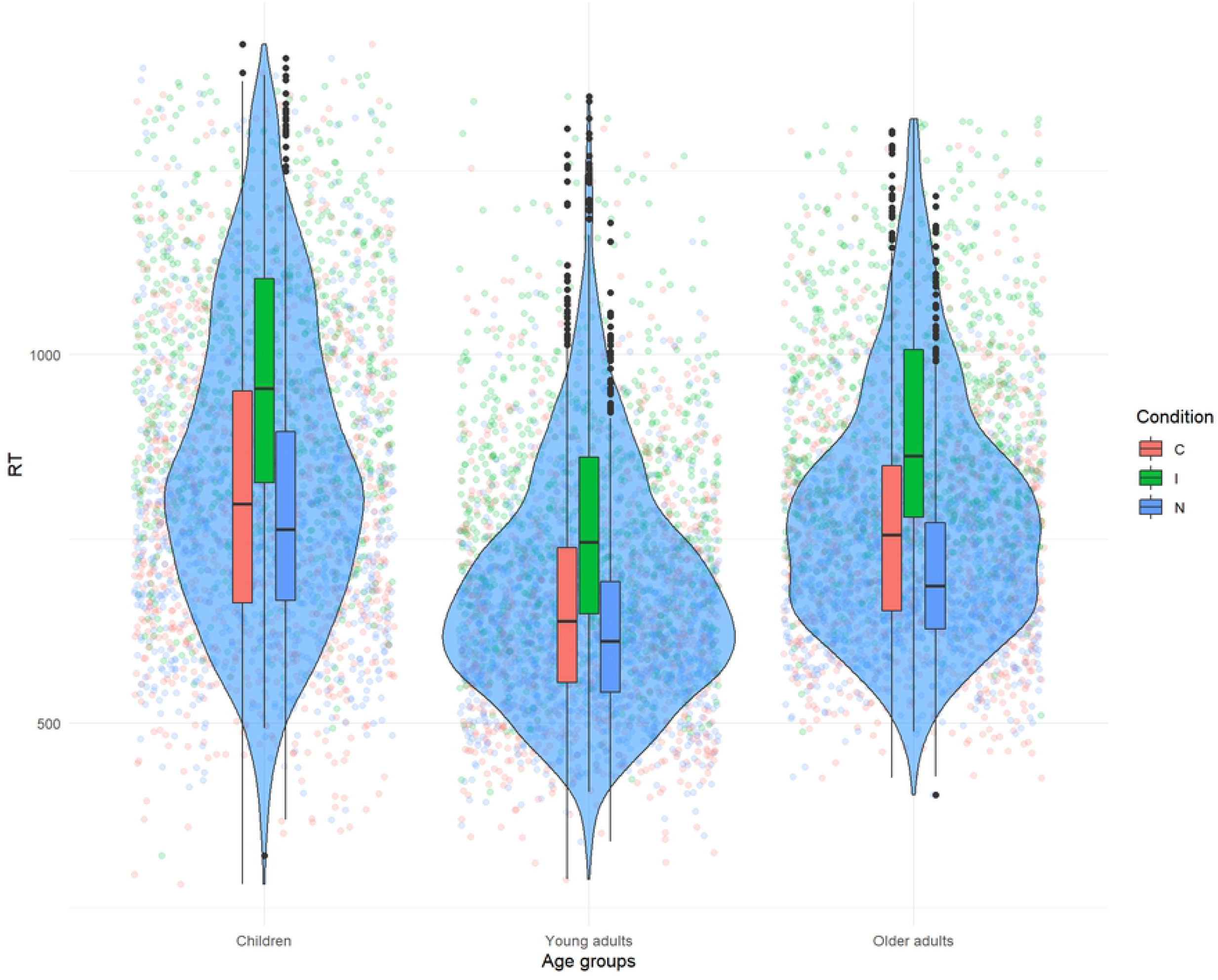
distribution of the latencies per conditions and age groups.

As a reminder, the linear mixed model included the conditions and the age groups as fixed effects, and the subjects ID variable as random intercept. The model highlights a significant main effect of condition on reaction times (F(2,18.9) = 111.68; *p* < 0.001), a significant main effect of age groups (F(2,50) = 10.74; *p* < 0.001), and a significant interaction between the two (F(4, 8900.7) = 13.56; *p* < 0.001). The decomposition of the interaction reveals that children (t(132.6) = −8.83; p<0.001), young (t(125.27) = - 6.04; p <0.001) and older adults (t(131.73) = −8.54; p<0.001) show significantly increased latencies in the incongruent condition. As emphasized by the estimates of the model, the results show a modulation of the effect’s magnitude among conditions and age groups, suggesting that the interference effect is the strongest for children (estimate = 104.96), while it is the weakest among the young adults group (estimate = 35.98), and intermediate for older adults (estimate = 94.86).

Regarding accuracy, the results highlight a significant main effect of conditions (χ^2^(2)=162.44; *p*<0.001), a significant main effect of age groups (χ^2^(2)=50.43; *p*<0.001) but no interaction between the two (χ^2^(4)=4.8; *p*=0.3). The decomposition of the main effect of conditions suggests that congruent trials were less prone to errors than incongruent trials (Z = 6.68; p<0.001), but none of the other comparisons reached significance. The age groups effect shows that accuracy was significantly lower in children compared to young adults (Z = −4.52; p < 0.001), however no significant differences were observed between children and older adults (Z = −0.02; p ≈ 1) nor between young and older adults (Z = −0.02; p ≈ 1).

### 3.2. ERP results

**Figure 2:**
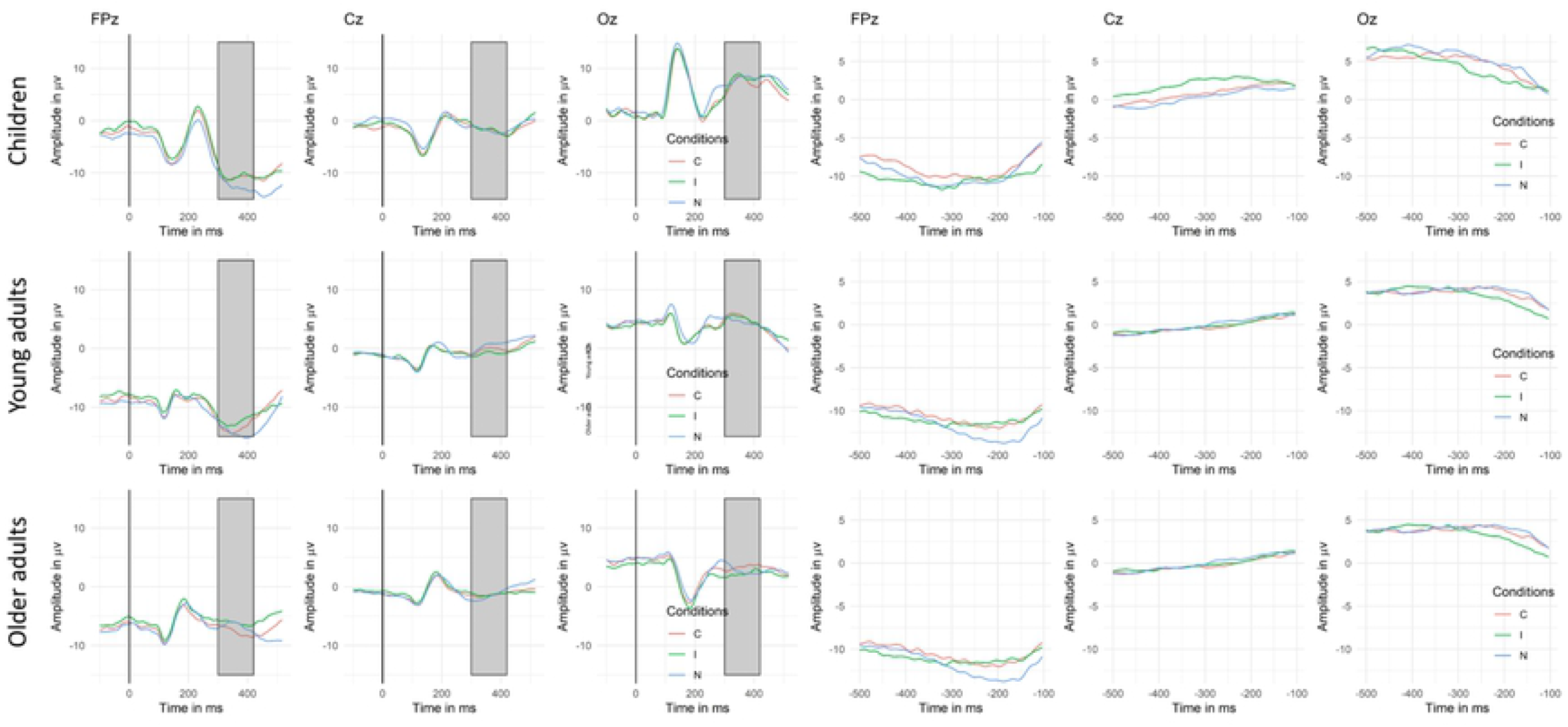
Exemplar waveforms of the grand averages of stimulus- and response-aligned ERP signal for the three conditions (C = congruent; I = incongruent; N = neutral) in each age group for the FPz, Cz and Oz electrodes. The peak analysis was also performed on the Fz and Pz electrodes. The gray rectangles shows the time window considered in the peak analysis (300 – 420ms).

#### 3.2.1. Mass univariate test and peak analysis

The results of the mass univariate tests on the waveform amplitudes run separately on the stimulus- and response-aligned ERPs are presented in Figure 3. Since neutral trials were very different from congruent and incongruent trials, it was suspected that most of the effect would be carried by the neutral condition compared to the other two conditions. That is what was observed since significant difference in amplitudes appeared on virtually the entire time window for both main effects of conditions and age groups as well as for the interaction as presented in Appendix S1. Since the present study focused mainly on the Stroop effect, contrasts including neutral conditions were discarded, and in Figure 3, only the contrasts between the incongruent and congruent condition are displayed for each age group.

The results of the mass univariate statistic highlighted different time windows of significant variations in amplitudes between congruent and incongruent trials in the three age groups (displayed in gray on Figure 3). First, the older adult group is the only one yielding significant differences between the two conditions in the stimulus-aligned analysis in the time-window from 260 to 520ms and on large clusters of electrodes. On the response-aligned analysis, children and young adults show significant effects of conditions in a similar time window (from −500 to −360ms), although not on the same electrodes, and with an additional time window (from −350 to −180ms) in children. Older adults presented a time window of significance closer to production (from −210 to −100ms).

The temporal window of interest of the N400 component did not show any significant effect for the children and young adult groups on the mass univariate tests. To ensure that our results were comparable to the rest of the literature, we conducted a peak analysis on a temporal window compatible with this component and showing the maximum difference based on a visual inspection. Repeated measures ANOVAs were conducted on an averaged time window ranging from 300 to 420ms (as shown on Figure 2) for the FPz, Fz, Cz, Pz and Oz electrodes. The models from all five electrodes are detailed in the Appendix (Appendix S2) but only the significant results on FPz are presented here. First, the model highlighted a significant main effect of conditions (F(2,100) = 6.94; *p* = 0.001), a significant main effect of age groups (F(2,50) = 4.45; *p* = 0.017) and a significant interaction between conditions and age groups (F(4,100) = 4.75; *p* = 0.001). Post-hoc decomposition achieved by a Tukey test showed that young adults present a significantly more negative deflection in the congruent condition compared to the incongruent one (t(100) = −2.45; *p* = 0.042), while this results was marginally significant in the older adults group (t(100) = −2.26; p = 0.066). Complete decomposition of the differences between conditions among the three age groups are presented un Appendix S3.

#### 3.2.2 Topographic analyses of variance (tANOVA)

The results of the topographic analysis of variance (tANOVA) displayed in Figure 3 showed different patterns across groups but were globally concordant with the mass univariate test. Regarding the stimulus-aligned signal, in children, a small time window appeared significant during the baseline, close to stimulus onset. In young adults, two small time periods suggested a change in topography, occurring at about 400 and 520ms. In older adults, several periods of significance appeared around 400ms post-stimulus, but also briefly during the baseline. In the response aligned signal, the same pattern as the mass univariate results were highlighted for children and young adults, namely two periods of significance for children and one, far from the response onset, for the young adults. However, older adults showed a discrepancy between the mass univariate statistic and the waveforms results. Indeed, the tANOVA shows a significant time window far from the response onset, while the mass univariate reveals an effect close to the onset of the response.

**Figure 3:**
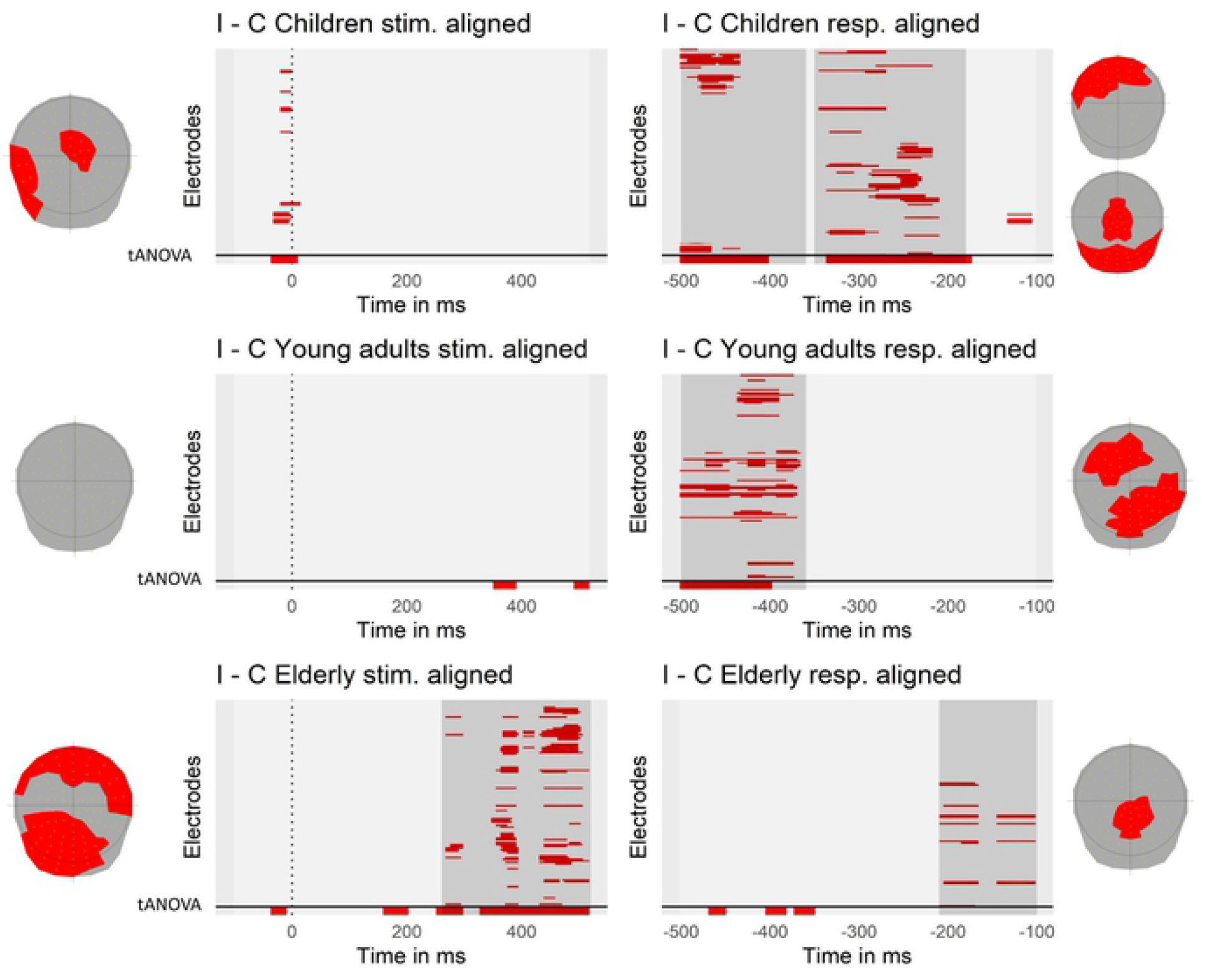
Significant differences in the contrast between congruent and incongruent conditions on each electrode and time-point for stimulus- and response-aligned ERPs within each age-group. The dotted vertical line represents the onset of the stimulus and the signal under the horizontal solid line at the bottom of each plot represents the tANOVA results specific to each contrast. Periods of consecutive significance shorter than 4 time frames (~16ms) were discarded. Clusters of significance are highlighted in gray. The topographies on the side represent the spatial distribution of the significant electrodes constituting the different clusters.

#### 3.2.2. Microstates analyses: segmentation and characteristics of the maps

The TAAHC algorithm identified five microstates explaining the signal on the stimulus-aligned (Maps 1 to 5) and four microstates for the response-aligned signal (Maps 6 to 9). Descriptive statistics about duration of the maps can be found in Table 2.

**Table 2:**
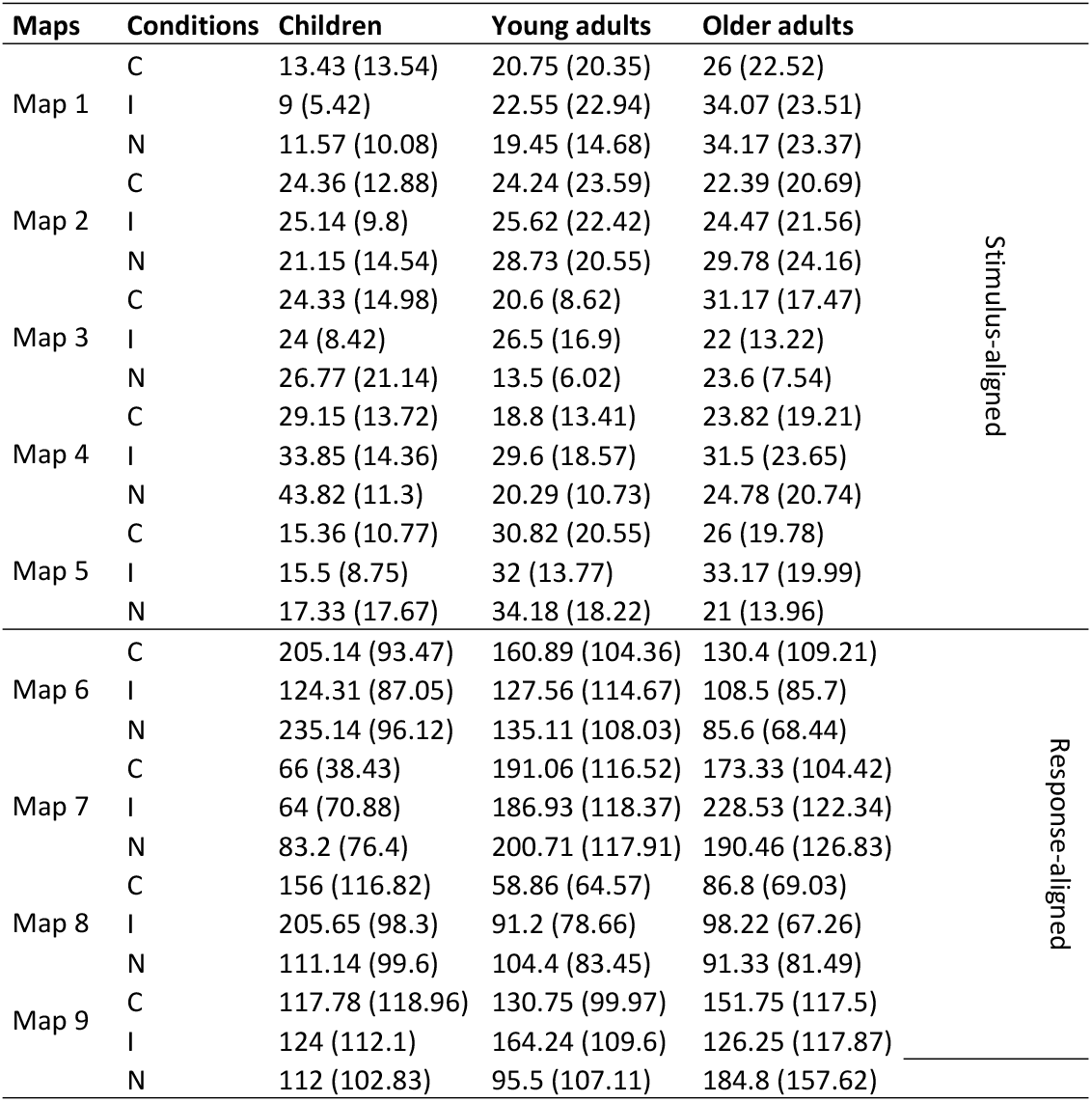
mean and standard deviation of the duration of each map across conditions and age groups.

#### 3.2.3. Microstates analyses: presence or absence of the template maps

The first analyses were performed on the presence of each map in the individual averaged signal per condition. For detailed presence information of each microstate map, see Appendix S4. Since the aim was to assess whether the maps explained the signal among the different age groups or conditions, the model tested for an interaction between conditions and maps as well as between maps and age groups. Results showed a significant main effect of maps (stimulus-aligned: χ^2^ = 39.2; p <0.001; response-aligned: χ^2^ = 13.08; p = 0.004), and a significant interaction between the maps and the age groups (stimulus-aligned: χ^2^ = 79.69; p <0.001; response-aligned: χ^2^ = 84.15; p <0.001). The other effects did not reach significance. Detailed decomposition was achieved by a Tukey test. The most relevant comparisons showed that some significant differences were observed on the stimulus-aligned signal among the three groups. Maps 1 and 2 were more present in the signal of older adults than in children (respectively: z = −2.91; p = 0.01 and z = −2.37; p = 0.047). Map 3 was more specific to children (older adults – children: z = 4.91; p <0.001; young adults – children: z = 4.89; p <0.001), while map 5 was more present in young adults, compared to older adults (z = −3.87; p <0.001). Regarding the response-aligned signal, results consistently show that map presence differs drastically among children and adults. Indeed, only comparisons involving children and adults returned significant while none of the comparisons between the two adult groups did. As presented in Figure 4, maps 6 (Children – Older adults: z = 2.73; p = 0.017; Children – Young adults: z = 2.81; p = 0.014) and 8 (Children – Older adults: z = 3.28; p = 0.003; Children – Young adults: z = 4.44; p <0.001) were significantly more characteristic of children while maps 7 (Children – Older adults: z = −2.86; p = 0.012; Children – Young adults: z = – 4.06; p <0.001) and 9 (Children – Older adults: z = −3.01; p = 0.007; Children – Young adults: z = −3.5; p = 0.001) were more representative of adults, young and older. As a consequence of these results on the presence of maps across groups, only maps present on a sufficient number of subjects within a group were compared across conditions. Even though all data were still included in the statistical model at all time, for each comparison among the groups and/or conditions, only the comparisons among the maps present in Figure 4 were interpreted.

#### 3.2.4. Microstates analyses: duration of the microstates

The statistical model on microstates duration (Hurdle model) in the individual ERPs included as fixed factor the maps labels, the age-groups, as well as the conditions of the Stroop task. All interactions were entered in the model. The random structure included the subjects’ ID as random intercept, and post-hoc decomposition of the effects was achieved by Tukey tests. Median durations of the microstates maps per age group are presented in Appendix S5

As shown in Table 3, results of the conditional model (the zero-inflated part will not be interpreted) on both stimulus- and response-aligned signal suggest a significant main effect of maps and conditions, no main effects of age groups, but significant interactions between the maps and conditions, a significant interaction between maps and age groups and a significant interaction between conditions and age groups. Finally, a triple interaction between conditions, age groups and maps appeared significant as well.

**Table 3:** results table presenting the conditional part of the zero-truncated Poisson generalized linear mixed models performed on the microstates duration.

|  | Stimulus-aligned |  |  | Response-aligned |  |  |
| --- | --- | --- | --- | --- | --- | --- |
|  | Chisq | Df | Pr(>Chisq) | Chisq | Df | Pr(>Chisq) |
| Maps | 218.36 | 4 | <0.001 | 348.7 | 3 | <0.001 |
| Conditions | 33.22 | 2 | <0.001 | 6.33 | 2 | 0.042 |
| Age group | 3.21 | 2 | 0.201 | 0.1 | 2 | 0.953 |
| Maps*conditions | 40.44 | 8 | <0.001 | 150.45 | 6 | <0.001 |
| Maps*age groups | 460.69 | 8 | <0.001 | 627.93 | 6 | <0.001 |
| Conditions*age groups | 31.92 | 4 | <0.001 | 12.72 | 4 | 0.013 |
| Maps*conditions*age groups | 137.01 | 16 | <0.001 | 254.54 | 12 | <0.001 |

**Figure 4:**
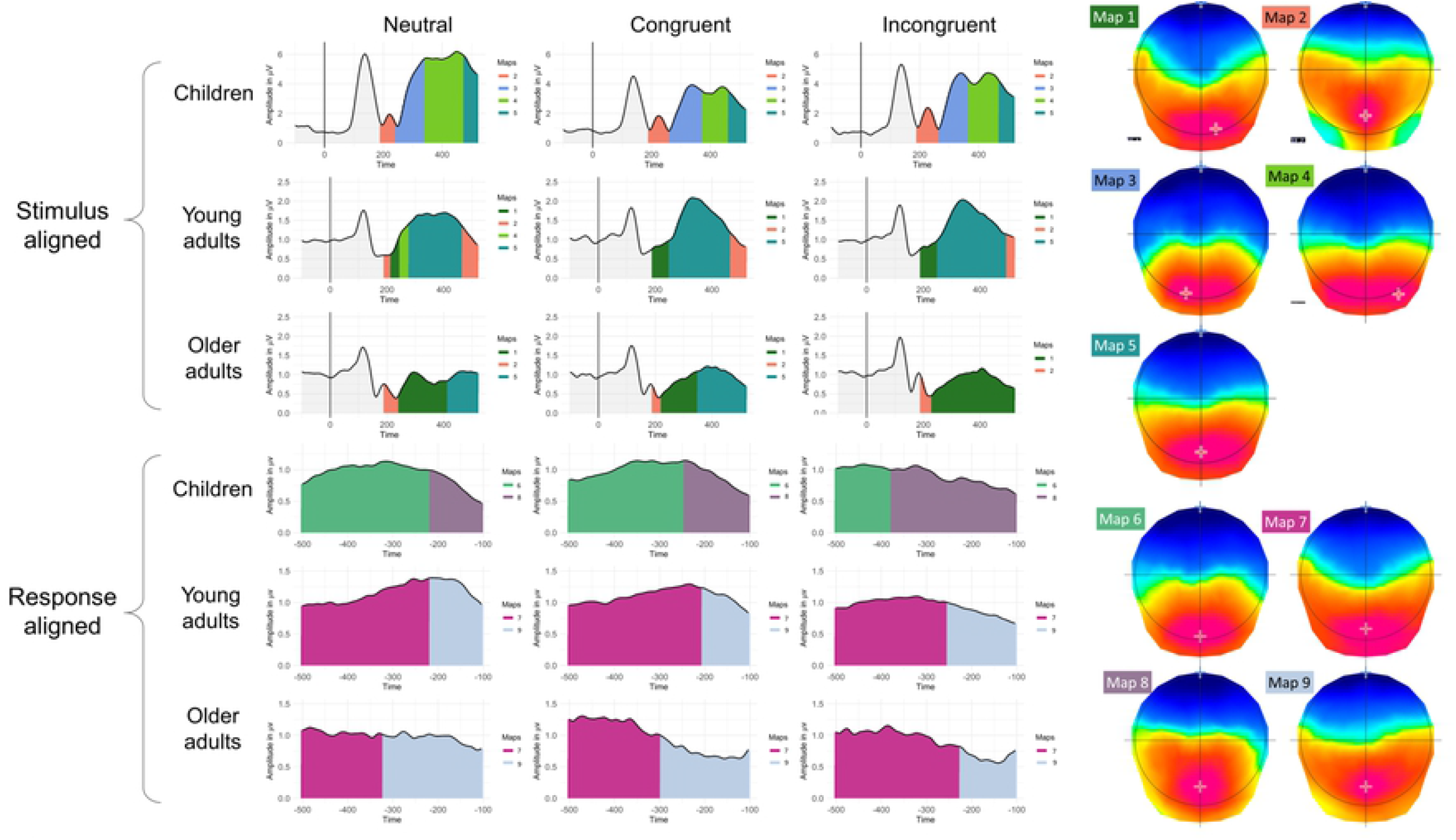
Periods of stability of the electric field at the scalp are represented under the GFP curve of the grand average ERPs with color codes for stimulus-and-response-aligned signal, for each age group and condition. The corresponding template maps are displayed on a continuum from red (positive) to blue codes (negative).

It is noteworthy to mention that the differences observed on the most distant map from the alignment point (i.e. microstates at the upper and lower borders of stimulus- and response-aligned data respectively: for children, Maps 5 and 6, for young adults, Maps 2 and 7, and for older adults, Maps 1 and 5 for stimulus-aligned and Map 7 for response-aligned signals) were discarded. Indeed, these maps are susceptible to carry a bias due to differences in latencies among conditions and age groups whereas the analyzes were carried out on identical fixed periods. In other words, the duration of the most distant maps from the alignment point might be modulated by the fixed length of the epoch across conditions and age groups.

First the comparison between age groups was considered (disregarding the differences among the conditions) and in second time, the differences among the conditions will be reported.

As shown in the previous analysis on the presence/absence, larger differences are observed between children and adults than between the two adult groups for the stimulus-aligned signal. According to Figure 4, Map 1 can be compared between the two adult groups and only Maps 2 and 5 can be compared between the children and the two adult groups. Results suggest that Map 1 had a significantly longer duration in the older adults group compared to the young adults group (t(704) = 3.26; p = 0.003). Maps 2 and 5 did not show any significant difference between any of the groups. Furthermore, some maps differentiate the congruent and incongruent conditions. For the children’s group, only Map 4 was marginally significantly more present in the incongruent condition compared to the congruent one (t(704) = −2.07; p = 0.097). For the young adults group, no effect was reported, and concerning the older adults group, Map1 (t(704) = −3.95; p <0.001) and Map 5 (t(704) = −3.12; p = 0.005) were significantly more present in the incongruent condition. Nevertheless, it is worthy to mention that on the segmentation depicted on Figure 4, Map 5 seems much more present in the congruent condition. This might be related to the fact that, as presented in Figure 4 and above in the absence/presence analysis, only 50% of older adults showed this map in the incongruent condition.

Regarding the response-aligned signal, only the two adult groups can be compared on Map 9. The results show a longer microstate for the older adults for the neutral condition (t(563) = 5.25; p <0.001) while the same map was marginally significantly less represented in older adults regarding the incongruent condition (t(563) = −2.35; p = 0.05). Moreover, for children, Map 8 lasted significantly longer in the incongruent condition compared to the congruent one (t(563) = −5.34; p <0.001). For young adults Map 9 lasted significantly longer in the incongruent condition compared to the congruent one (t(563) = −3.65; p <0.001), while the effect for older adults was only marginally significant (t(563) = 2.34; p = 0.051).

#### 3.2.5 Microstates: analyses on the onset of the microstates

To confirm that differences in duration explained differences among the groups, the onset time of each microstate was analyzed for each subject. The model included all main effects and interactions among maps, conditions and age groups. For this analysis, maps at the extremity of the temporal window have been considered.

Regarding the stimulus-aligned data, the results completed the ones observed on the duration of the maps and highlighted a significant main effect of maps (χ^2^ = 230.25; p <0.001), a significant main effect of age group (χ^2^ = 11.24; p = 0.004), and a significant interaction between maps and age groups (χ^2^ = 175.58; p <0.001). As stated above, only relevant post-hoc comparisons were interpreted, namely differences among all age groups for Maps 2 and 5, and also Map 1 for the adult groups. Map 2 had a significantly earlier onset for children (z = −5.57; p <0.001) and older adults (z = −5.26; p <0.001) compared to young adults. Regarding Map 5, as presented in Figure 4, this map had a significantly delayed onset compared to both young (z = 6.04; p <0.001) and older (z = 4.77; p <0.001) adult groups. Regarding Map 1, older adults have a significantly later onset compared to young adults (z = 2.86; p = 0.012).

Regarding the response-aligned model, the results point out a significant effect of maps (χ^2^ = 101.22; p <0.001), a significant interaction between maps and age groups (χ^2^ = 45.64; p <0.001), as well as a marginally significant interaction between maps and conditions (χ^2^ = 11.27; p = 0.08). Since the triple interaction including age groups was not significant, this interaction was not decomposed. Post-hoc decomposition of the interaction between Maps, age groups and conditions did not highlight any relevant significant effect. Indeed, the only relevant comparison among the groups was between young and older adults on Map 9 according to the segmentation presented on Figure 4, which failed to reach significance (z = −1.29; p = 0.403).

#### 3.2.6 Source localization of the microstates

To better understand the brain networks involved in the microstate maps, a source localization procedure was performed on the grand averages, based on the timing of the microstates maps bearing the differences among groups or conditions (Figure 5). In the stimulus-aligned signal, Map 3 appeared to be specific to the children group but did not show a significant difference between the congruent and incongruent conditions. This map reflects mainly activations in the bilateral (but mainly right) inferior temporo-parietal regions. In the adult groups, Map 5, which encompasses the time window of the N400, seems to carry the Stroop effect in the older adults group but not in the young adults group. In young adults, congruent items elicit mostly parieto-occipital regions, while the maximum activation located in the ACC and fronto-mesial regions are reported in the incongruent condition. In the older adults group, the condition does not seem to impact the brain regions engaged. This age group displays a globally similar pattern as the one observed in young adults in the congruent condition.

Regarding the response aligned signal, in the last 300ms before the response onset, the three age groups show different patterns. Children and young adults show a strong fronto-mesial activation in the incongruent condition. In the congruent condition, young adults show a distributed left temporal network while children consistently show an activation of the fronto-mesial areas. Conversely to the other two age groups, older adults show a temporo-insular activation in both congruent and incongruent conditions.

**Figure 5:**
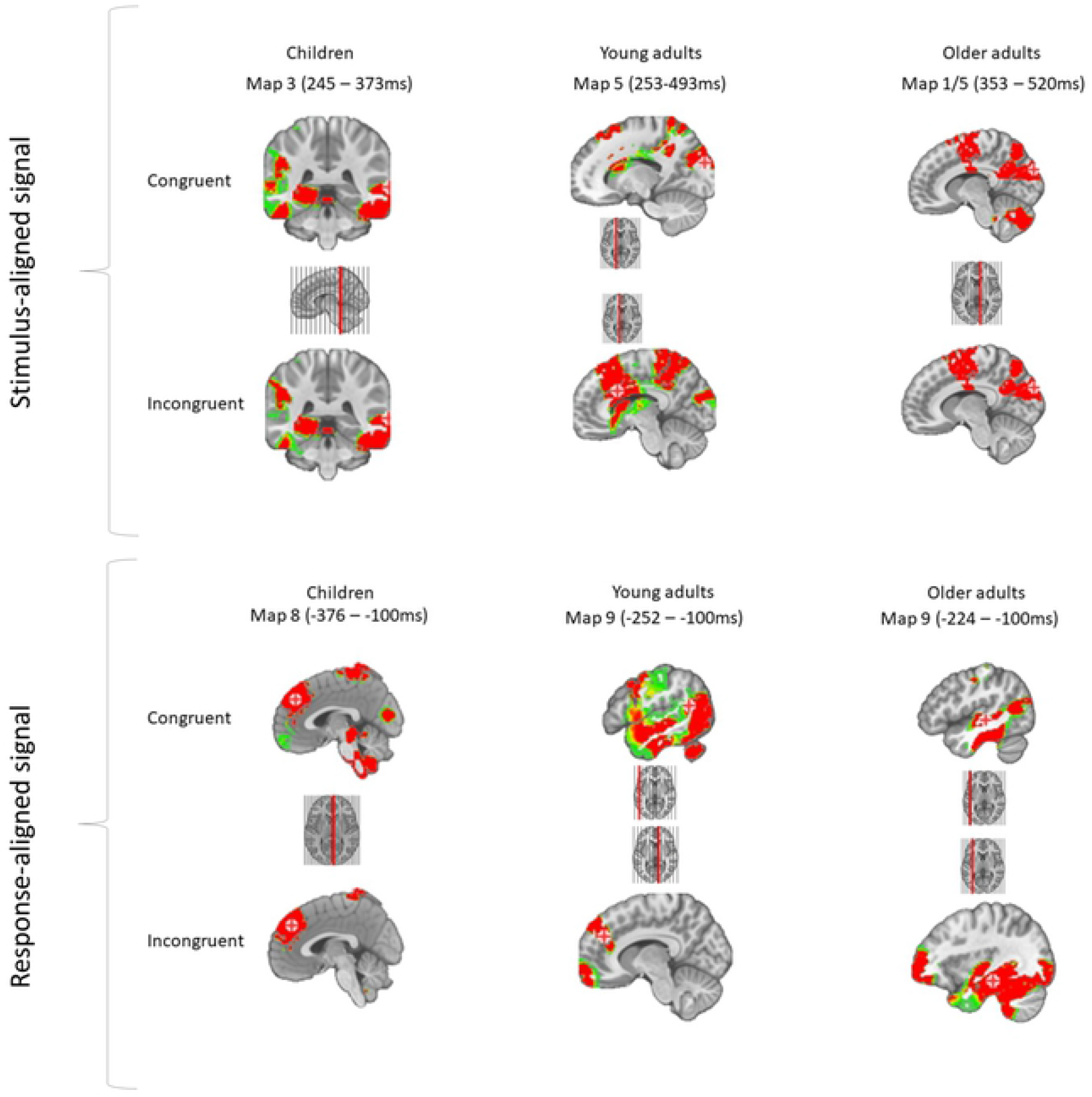
sources relative to the timing of the microstates disentangling groups or conditions in regards of the Stroop effect. Red cross indicates the maximum of activation.

## 4. Discussion

The present study aimed at understanding how the Stroop effect with its two levels of conflict (perception- and resolution-based conflicts) evolves with age, since latencies are longer for children and older adults. The present results confirmed the larger latencies in both children and older adults reported in previous studies, but the ERP results indicate different underlying processes in development and aging. In the following, we will first briefly comment the behavioral results before digging separately the issues related to development and aging. Finally, we will discuss additional findings such as the robustness of the N400 and the reliability of the SP600 components before concluding.

### 4.1. Behavioral results

The behavioral results supported the expectations, showing a strong Stroop effect on the latencies for each group while the measurement of accuracy did not highlight clear effects. Indeed, even though the congruent condition was less prone to errors than the incongruent one, no modulation across the age groups was emphasized. Across the three conditions, children were the group showing the largest error rate. Regarding older adults, even though the difference was not significant, it is worthy to note that this group was more accurate compared to their young counterparts, which could translate a change in the speed-accuracy tradeoff between the two groups. Young adults might be more focused on rapidity to provide a swift response while older adults might rather favor accuracy. This might be the first argument claiming that aging is not maturation in reverse [27]. To explore more precisely how maturation and aging differ, microstates analyses were performed on the task to help going beyond the limitations of behavioral and classical ERP methods.

### 4.2. From childhood to adulthood

#### 4.2.1. Conflict detection – stimulus-aligned signal

The signal of the three age groups includes the same electrophysiological components, namely the P100, the N1/N170, the P300 and a long lasting positivity usually labeled as the SP600. However, as shown in Figures 2 and 4, children tend to display two components separating the P300 from the SP600. Previous results from the literature have shown that school-aged children might show a delayed P2, peaking around 400ms [84]. Regarding the Stroop task, even though one study presented different components between children and adults [85], none of the other studies (all using pseudo or manual Stroop tasks) showed different components between age groups [33,35,86]. Since this particular pattern has been shown only in picture naming and by the results of the present study, it is most probably attributable to language processing.

Microstates allowed to segment the ERP signal in periods of quasi-stable electrical configurations reflecting activated brain networks at certain temporalities. We will first describe here the within-groups comparisons for children and young adults, and in a second time discuss the differences in the brain networks involved between the two groups on the stimulus-aligned signal, and then discuss response-aligned results. First, our data confirmed that in children and young adults groups, the same processes were involved in congruent and incongruent trials and only differed in duration. Such a result was already reported in the literature [16,57] with young adults and favors the idea of a single conflict detection mechanism activated continuously for all types of trials.

The Stroop effect (incongruent vs. congruent) was not significant on the duration of any of the microstates found on the children stimulus-aligned signal, however, a trend was highlighted regarding Map 4, which appeared on the N400 time window (367 – 470ms). It is worth to mention that there were no significant differences across conditions in this time window on the waveforms analyses nor the tANOVA. In accordance with this tendential result, none of the maps showed significant differences between conditions in the young adults group. This absence of results is not in line with those by Khateb and colleagues but rather replicate those by Ruggeri et al [57], who also showed an elongation of a microstate map close to the onset of the production.

Despite similar processes across conditions in children, as already observed in young adults, it remains to be discussed if children are recruiting the same brain networks as young adults and if the timing of the shared microstates is different. When comparing the topographic maps in children and young adult groups, it was observed that Map 3 (263 – 362ms) was specific to children. This Map 3 appears following the N170 component, which seems common across groups (Map 2). However, the time period of Map 3 is not yielding the interference effect since the map was not significantly different between the congruent and incongruent conditions. Seemingly, the process involved in the P300 component is still immature in children. Here, it may be related to reading [87] or to lexical selection [88] processes, or both. The spatial localization of signal in this time window showed an activation in the parieto-occipital regions, and especially in the right hemisphere, which have been related to the processing of a word compared to a fixation cross [89,90], as well as words reading compared non-words [90], rather suggesting that Map 3 represents word reading processes in children. Alternatively, a change in the same temporality has been proposed in the aging literature, stating that the P300 was modulated between young and older adults [12,17,48]. As described below, the authors attributed this modulation to a preparation component related to the attentional processes aiming at facilitating the processing of the conflict.

In other words, on the stimulus-aligned signal, children and adults showed globally the same maps except for an additional one in an early time window. Now, regarding the duration of each of these common processes, Map 5 was delayed in children (475 – 520ms) compared to young adults (265 – 477ms). This difference is compatible with the results described above suggesting the involvement of an additional map in the children’s group. The delay is probably the consequence of this additional activation (see Figure 4). Interestingly, this map is also the one present in the time window of the N400 component for the young adults, supporting the hypothesis that children show immature specific brain networks involved in conflict detection instead of a lack of maturation of the attentional network, slowing down each process.

#### 4.2.2. Conflict resolution – response-aligned signal

The results on the response-aligned signal are much clearer than on the stimulus-aligned data. First, once again, the results suggest that the brain processes involved within groups in the congruent and incongruent conditions are identical and only differ in duration between the two conditions. Indeed, for the children’s group a Stroop effect was highlighted on Map 8, which appeared in the signal close to the response onset (−279 – −100ms) and a similar finding was reported in young adults on Map 9 (−224 – −100ms). In other words, both groups showed a Stroop effect globally at the same temporality but on different topographic maps. According to the source localization analysis, children show a much more reliable activation of the ACC compared to the other two groups who tend to use more strongly their left temporal lobe to process Stroop items. Nevertheless, it appears that the maximum activation for young adults is located in the ACC for the incongruent condition (as for children), but in the temporal lobe for the congruent condition despite the absence of significant differences in the topographies. This discrepancy might be explained by the fact that during this time window, left-temporal structures as well as the ACC were activated but only the maximum activation switched from one region to the other. This finding implies that the conflict resolution processes in children are not yet identical to those of young adults (different brain networks involved) but since the timing of the maps that encompass the interference effect is comparable, these mechanisms are probably the immature version of the adult ones and no additional process is needed to resolve the conflict in children. This interpretation is also in line with developmental studies showing that the SP600 is still maturing after the age of 12 [33,34], since different brain networks were involved in this time period.

### 4.3. Aging

Even though from the point of view of latencies older adults tend to show the same increased response time regarding the interference effect as children, the precise mechanisms involved remain to be explored. Children showed underdeveloped specific brain networks but older adults might show a different pattern such as similar networks differing only in duration, then favoring the general slowing hypothesis.

#### 4.3.1 Conflict detection – stimulus-aligned signal

Once again, the results favor the hypothesis of identical mechanisms involved in congruent and incongruent conditions, already discussed above. Moreover, older adults showed a longer lasting period of electrophysiological stability at the scalp corresponding to Map 1 in the incongruent condition (236 – 520ms) compared to the congruent one (228 – 352ms), carrying the Stroop effect. Interestingly, this map starts in the temporal window of the P300 component, which has been shown to be modulated in aging for the Stroop task [12,17,48]. Nevertheless, the topographical analysis add two new pieces of information compared to the previously known conclusions of the peak analysis on the P300. First, the results show that the brain networks underlying this component are partially different in young and older adults (Map 5 for young adults and Map 1 for older adults). Second, Map 1 (C: 228 – 352ms; I: 236 – 520ms) lasted significantly longer in the incongruent condition compared to the congruent condition, which suggests that in older adults the conflict at the perception level is more widely represented and starts earlier in the signal compared to young adults. The sources underlying the time-window of maps 1 and 5 suggest the different generators involved in both adults group, showing that the ACC is more reactive in young adults than in the older adults group. In any case, these results have to be nuanced regarding the N170 component (negative component peaking at ~180ms with strong posterior negativity and centro-fronal positivity). Indeed, the results showed that young adults differed from the other two groups on the microstate present at the timing of this component. The absence of this component for the young adult group (as indicated by the GFP curve on Figure 4) seems to show that young adults directly activate the network responsible for conflict detection. This finding is in line with literature showing that the N170 might be linked to the preparation to resolve the conflict [10,15], which is translated by an earlier onset of the Map 1 in young adults compared to older adults.

The last point to be discussed is the difference in involvement of the brain networks among the different maps between young and older adults. The results showed that globally the same topographies were involved in both adult groups, except for Map 5 (young adults: 265 – 477ms; older adults: 388 – 520ms) which tended to be more present in the young adults group. These results could be explained by two different hypotheses and have to be related with the extreme duration of Map 1 in the older adults group. It is possible that, the duration of Map 1 pushed Map 5 outside the temporal window of interest. An alternative explanation would be that Map 5 was suppressed to compensate the wide duration of Map 1. In other words, the processes involved in older adults take longer, at least for the conflict detection processes without the involvement of compensatory mechanisms, which supports the general slowing hypothesis. To claim that this hypothesis explains best the results, we need to confirm that microstates on the response-aligned signal follow the same trend.

#### 4.3.2. Conflict resolution – response-aligned signal

In the response-aligned signal the data also showed the same microstates involved in congruent and incongruent conditions. Even though the effect reached significance in the young adults group while being marginally significant in the older adults group, Map 9 (young adults C: −204 – −104; young adults I: −252 – −104; older adults C: −296 – −104; older adults I: −224 – −104), observed around 150ms before the vocal onset, seems to play a role in the response-based conflict processing of both adult groups. The weaker effect in older adults compared to young adults might be attributed to a higher cognitive load level already for congruent items, hence reducing the difference between congruent and incongruent trials. This could imply that simply dealing with two stimuli dimensions (color font and color word) is interferent without needing the two dimensions to be contradictory. When considering results of the source localization analysis, it appears that older adults activate strongly the left temporal areas to process congruent and incongruent items, while young adults seem to rely more on the frontal regions, despite similar topographies observed in the two groups for this time window. Here again, the same brain network is activated but only the maximum activation changes from one condition to another.

To conclude on aging, both stimulus- and response-aligned signals showed that the two adult groups displayed globally similar maps however differing in duration, which fits best the general slowing hypothesis. As a consequence, the less efficient performances observed in aging are rather to be attributed to a decline in attentional resources than in a specific decline in executive functions.

### 4.4. Integration of results on development and aging

To sum up, the present results led us to conclude that during development, the brain mechanisms involved are different than those observed in young adults. An additional network is required to process conflict detection and the other mechanisms seem to be involved differently. In aging, the data told a different story, favoring the general slowing hypothesis. Even though the two groups show radically different patterns, some elements can be relevant for both maturation and decline. On the response-aligned signal, all three groups showed an interference effect (incongruent vs. congruent condition) on the last brain network activated before the onset of production. This microstates however underlies different sources for the three groups. It appears that children rely mostly on the ACC to solve the conflict and even process congruent items, while young adults activate strongly this structure only in incongruent situations along with older adults using mostly their left temporal lobe. When considering the three groups as a continuum, it might be possible that with age, the interference processing uses more semantic features than executive functions, as suggested by Spreng and Turner [91]. They hypothesized that since at first, children can rely only on fluid intelligence and since in aging, adults tend to show a higher crystallized intelligence but lower fluid intelligence abilities, architectural changes along the lifespan experience a transition from fluid to crystallized intelligence engaged to perform cognitive tasks. This theory could explain the changes observed in the source localization analysis, however interference effect (incongruent vs. congruent trials) within each age group remains to be discussed. The interference effect present in all age groups in the response-aligned ERPs can be discussed in the framework of two different cognitive models, namely general models of language production [88,92] as well as the horse race model [93]. In neuropsycholinguistic models, the time window close to production has been associated with the phonetic encoding or motor speech planning, i.e. the preparation of the articulatory scores, encoding the motor pattern to produce the response or with self-monitoring processes [94]. From this point of view, the Stroop effect is compatible with a response-based conflict model but the linguistic stage impacted by this interference effect is rather speech encoding than language. According to these same models, this timing of activation might be related to the self-monitoring of the response [95,96]. Since models on language production and of the Stroop task evolved in parallel with very few interactions, the monitoring processes engaged close to the onset of the production and described in the general model on language production could reflect the same mechanisms as those involved in the conflict resolution of the Stroop. This interpretation is also in line with the response exclusion hypothesis [97,98]. According to the authors, the closer the distractor from the target semantically, the smaller the interference effect. These results were interpreted as an argument against the concept of competition in the lexical selection process, implying that the speaker suppresses the distractor (thus favoring the production of the target) during the post-lexical pre-articulatory stages of the word production model. The horse race model of Dunbar and MacLeod (1984) is now often used to describe the history of the cognitive models explaining the Stroop effect rather than to build new theoretical concepts on it. This model describes the cognitive processes involved in the Stroop task as two different processes, specifically color naming and word reading, triggered at the same time and competing only when reaching the response production stage. Nowadays, it has been shown that both processes interact more than expected by the model in early processing stages, although this model describes our data quite well.

Even though children and older adults show an interference effect, it does not seem to be that robust compared to response conflict. Moreover, the data suggest a strong and reliable response conflict which corroborates the predictions made by this model.

Even though the present results bring some clarification on the evolution of the Stroop effect, several issues remain open. Undeniably, it is still difficult to attribute a precise process to each of the brain networks highlighted by the microstates analyses. Future studies might need to replicate the result and compare tasks with different involvement of attentional abilities and of cognitive control. It seems also important to better understand the difference between verbal and manual tasks and how it relates to interference. Since differences were observed on the last stages of verbal responses, the divergent results with manual ones could be exploited in a direct comparison to clarify the underlying mental processes. Finally, some unexpected results were observed such as the absence of any Stroop effect on the N400 time window for the young adults group. Future studies need to clarify the reasons why this effect might not be as robust as previously reported, but in the meanwhile, a few of these reasons will be described in the following section.

### 4.5. Robustness of the N400 effect and interpretability of the SP600

The ERP results on waveforms using a mass univariate test showed that the N400 effect was not robust since it was only observed in the older adults group. When examining the waveforms presented in Figure 2 and given the results of the peak analysis performed on FPz, Fz, Cz, Pz, Oz electrodes, young and older adults show a significantly more negative deflection in the 300-420ms post-stimulus time window. Regarding children, a slightly more negative peak is observed around 420ms for incongruency on both electrodes even though later temporalities show a more negative trend for congruent than incongruent condition. The absence of a robust N400 effect may be due to the presence of neutral trials in the present study. Even though previous studies including neutral trials replicated the N400 Stroop effect [14,20,21], these studies usually used rows of “X” and manual responses. By varying the type of neutral trials, hence increasing the stimuli set size as well as the magnitude of the interference, the effect might have been reduced [99]. This, added to the strong correction applied by the mass univariate statistics might explain the disappearance of the effect. Moreover, for young adults, the tANOVA results tended to show a significant effect of condition on the time window of the N400 and children showed an interference effect on Map 4, present in the N400 time window. Another explanation might be related to the structure of the task. Since initially this task was designed to investigate the conflict adaptation effect [40], up to three repetitions of the same condition were allowed. Subjects might have benefit from this feature to anticipate the presence of a conflict in the next item. This point is supported by the results of Larson and colleagues [35] who analyzed the evolution of the conflict adaptation effect in development and did not report any significant N400 effect, but only a SP600 effect. Other pieces of evidence such as a small but significant effect in the mass univariate around stimulus presentation in the children signal or an early N400 effect in young adults bring some conviction to this interpretation.

Despite some discrepancies in the results relative to classical approaches, here this data-driven method allowed to describe more precisely the data, as stated below. Typically, ERP studies investigating the Stroop effect analyze a time window of about 1000ms following the stimulus onset, while subjects start responding on average around 682ms in our study (earlier with manual responses). For this reason, we preferred using two alignment points, one at the stimulus and the second at the response onset. This method allowed to overcome the response artifact on the signal, and its shift across conditions, which can contribute to the effect. Indeed, even though the SP600 effect was found when participants were asked to respond covertly [18], if the response is executed earlier in the congruent compared to the incongruent condition, the difference observed on the SP600 could be partly attributable to a shift in the motor activity related to the response. Nevertheless, aligning ERPs to the response has the inconvenience that the results cannot be precisely related to a SP600, but since the component is observed close to the response, it is very likely that a time window close to the response will correspond to the SP600 component. Interestingly, the mass univariate test did not highlight one single effect but rather three different time windows allowing to decompose the complexity of the effect.

## 5. Conclusion

The present study aimed at understanding if longer latencies in children or older adults compared to young adults could be attributable to the suboptimal functioning of a specific process, the addition of a compensatory process or a slowdown of all the involved processes. As discussed above, the results in children showed that the brain networks involved were substantially different from those engaged by the young adults group on both conflict detection and resolution. Children showed an additional microstate around 350ms and a Stroop effect carried by the brain network activated in the N400 time window, showing a more effortful conflict detection process for this group. Regarding conflict resolution, children tended to engage different microstates and an interference effect was again carried by a network activated right before the response onset. These results were interpreted as favoring the idea that the longer latencies compared to young adults could not be interpreted as a slowing of the same mechanisms, but rather the immaturity of specific processes related to conflict detection and resolution. Regarding aging, the results led to the opposite conclusion. Both stimulus- and response-aligned signals involve globally the same networks. However, they all seemed to be slowed down compared to young adults. These results favor the general slowing hypothesis, stating that the decrease in performance for an executive function task is not related to a specific decline of executive functions but rather a decrease in the attentional resources availability. It is noteworthy to mention that the results also highlighted that the N400 effect known to be characteristic of the detection of the Stroop conflict is not as robust as described. By using a classical peak analysis the results of princeps studies were replicated but vanished when using a more data-driven method. The SP600 component was however more robust but appeared to be more complex than described so far. Indeed, young adults and children showed a similar time window of the effect, while children presented a second time window of significance closer to the production of the response. Regarding older adults, conflict resolution effect was shown only later, in the last 200ms before production onset.

## Acknowledgements

We would like to thank the participants for giving their time for this study, as well as Tanja Atanasova, Raphaël Fargier and Giulia Krethlow who helped collecting the data. We also would like to thank Freda Ménétré for her opinions on the manuscript.

## Appendix

**S1:**
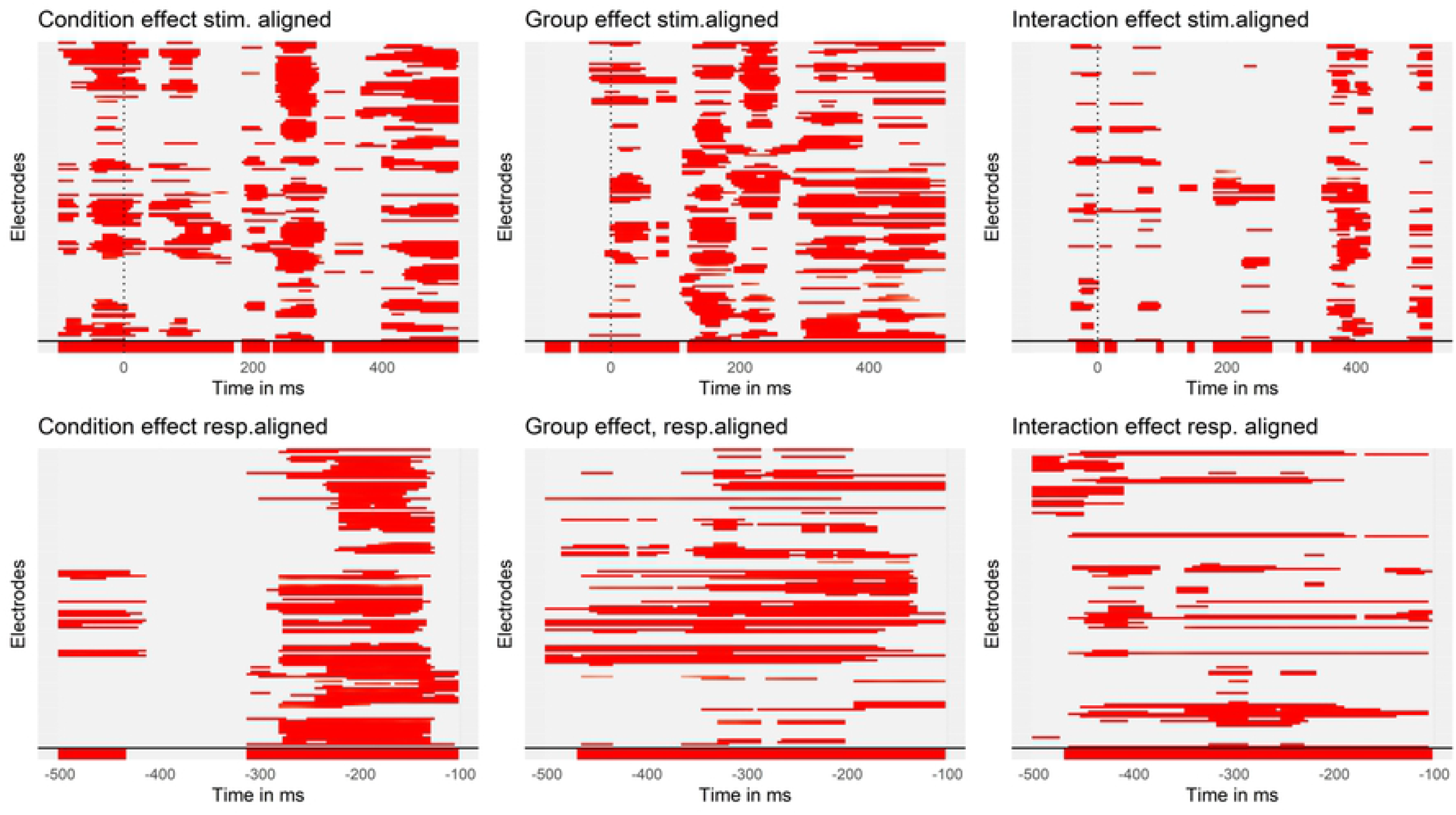
mass univariate tests on the waveform for each tested effect. The upper plots row represents the stimulus-aligned results while the bottom row represents the waveforms 500ms before the response onset. The last 100ms were counted as pre-articulation and were excluded from the analyses. The x axis of each graph represents the time in milliseconds and each electrode is represented on the y axis. The dotted vertical line represents the stimulus onset, and the signal beneath the solid horizontal line represents the tANOVA results specific to each effect.

**S2:** main effects and interactions of the repeated measures ANOVA models estimated in the context of the peak analysis.

| Electrodes | Effects | Sum Sq | Mean Sq | NumDF | DenDF | F value | Pr(>F) |
| --- | --- | --- | --- | --- | --- | --- | --- |
| FPz | condition | 71.76 | 35.88 | 2 | 100 | 6.94 | 0.002 |
|  | group | 46 | 23 | 2 | 50 | 4.45 | 0.017 |
|  | condition*group | 98.11 | 24.53 | 4 | 100 | 4.75 | 0.002 |
| Fz | condition | 3.53 | 1.76 | 2 | 100 | 0.95 | 0.39 |
|  | group | 29.66 | 14.83 | 2 | 50 | 7.99 | 0.001 |
|  | condition*group | 29.16 | 7.29 | 4 | 100 | 3.93 | 0.005 |
| Cz | condition | 0.68 | 0.34 | 2 | 100 | 0.68 | 0.509 |
|  | group | 4.63 | 2.31 | 2 | 50 | 4.6 | 0.015 |
|  | condition*group | 6.91 | 1.73 | 4 | 100 | 3.43 | 0.011 |
| Pz | condition | 2.27 | 1.14 | 2 | 100 | 1.51 | 0.227 |
|  | group | 4.09 | 2.04 | 2 | 50 | 2.71 | 0.077 |
|  | condition*group | 9.01 | 2.25 | 4 | 100 | 2.99 | 0.022 |
| Oz | condition | 1.7 | 0.85 | 2 | 100 | 0.34 | 0.711 |
|  | group | 17.64 | 8.82 | 2 | 50 | 3.54 | 0.036 |
|  | condition*group | 31.64 | 7.91 | 4 | 100 | 3.18 | 0.017 |

**S3:**
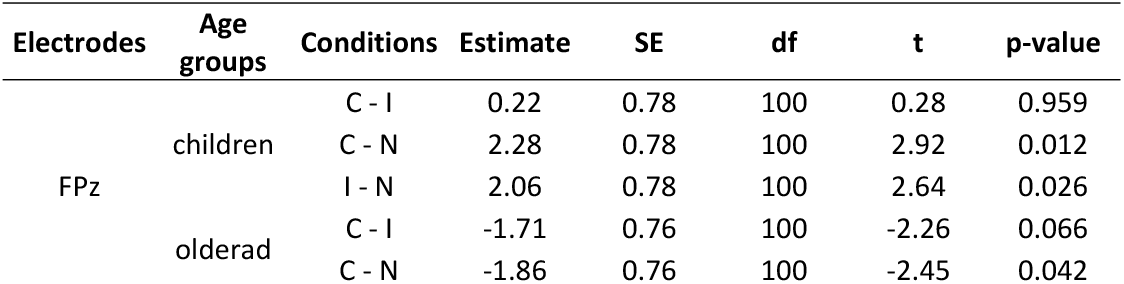
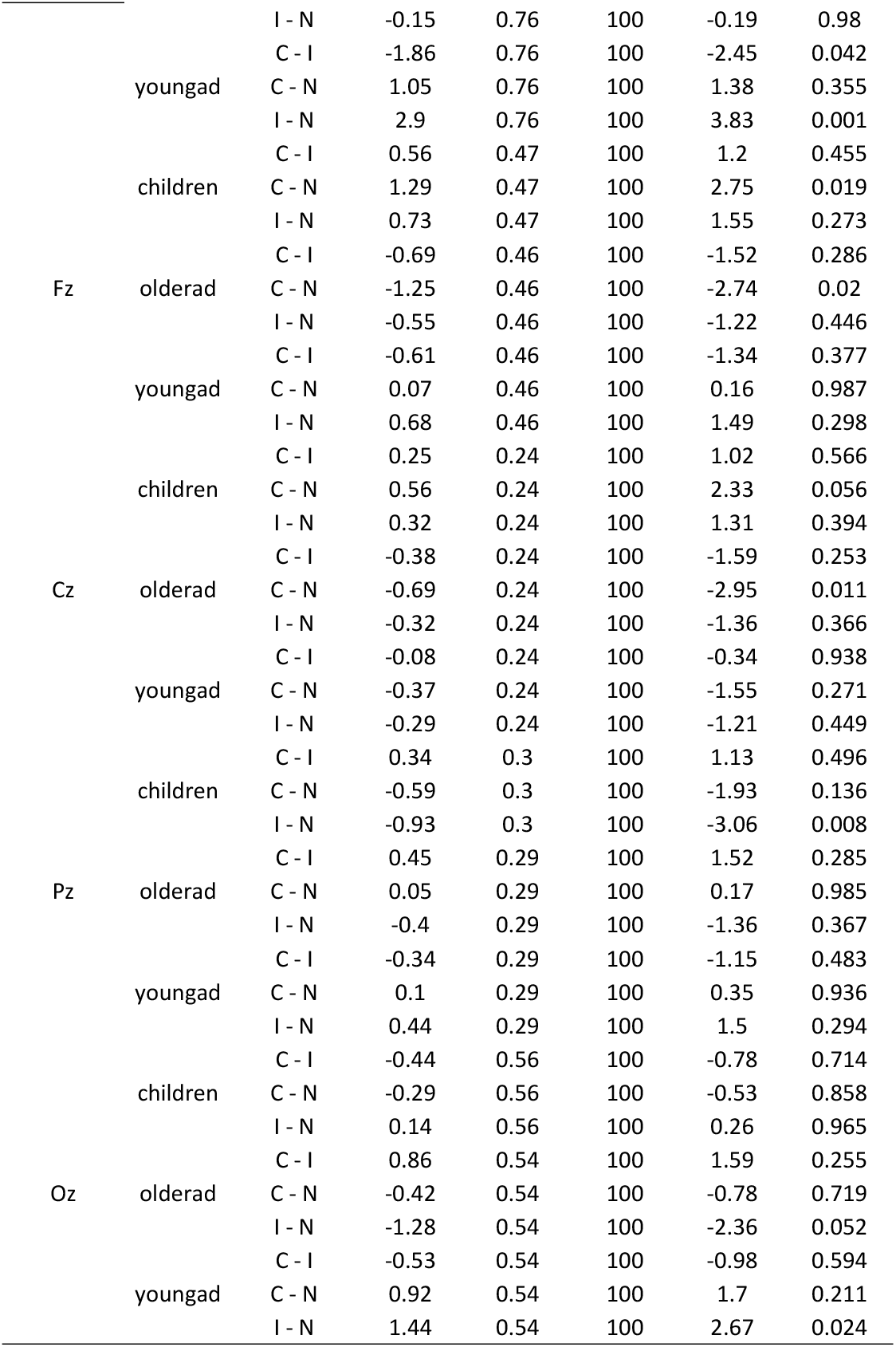
post-hoc decomposition of the interaction between conditions and age groups among the five electrodes retained for the peak analysis.

**S4:**
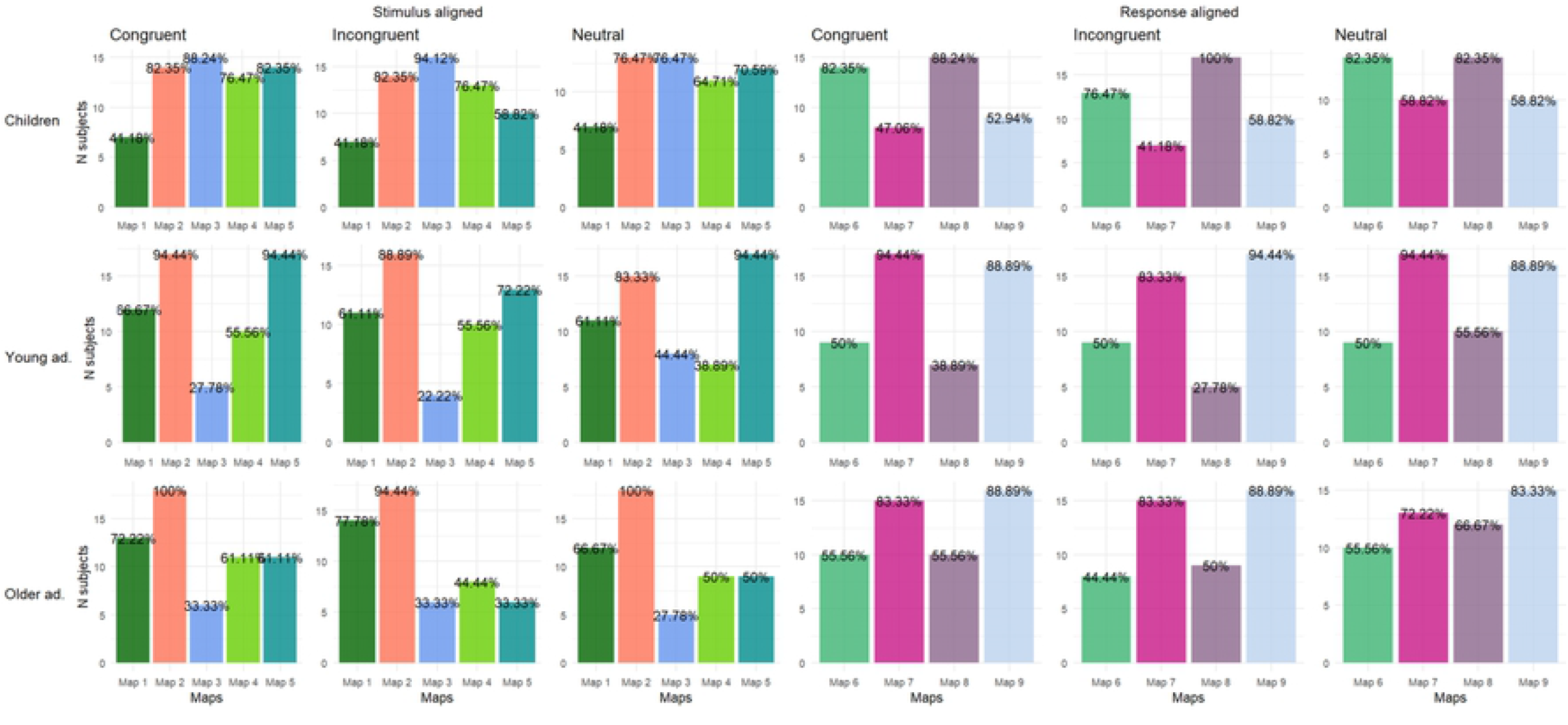
bar plot of the presence of the maps for each age group and condition. Values on the y axis represent the number of subjects in the age group for which this map was found, and the percentage represents the proportion of this number of subjects relatively to the number of subjects in the age group.

**S5:**
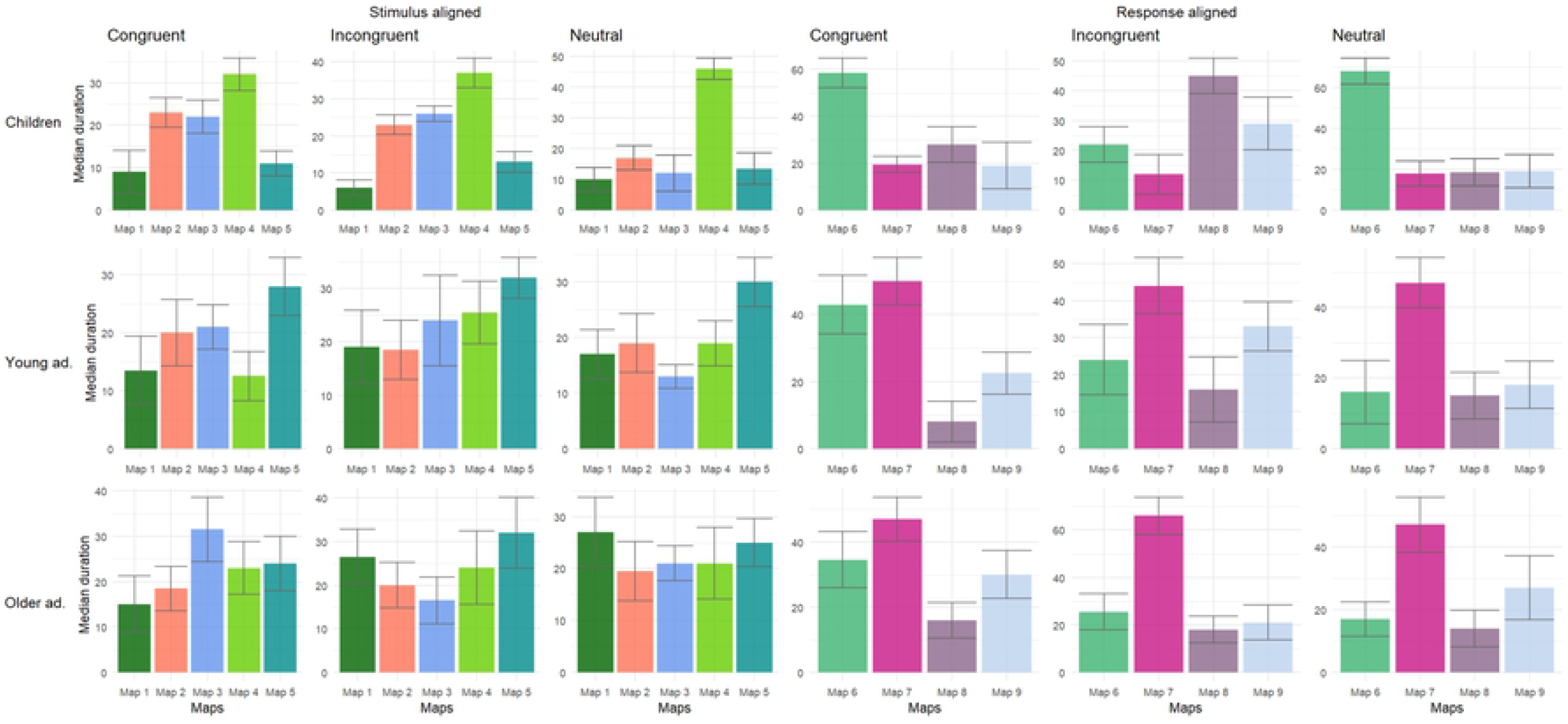
bar plots reporting the median duration (for subjects who showed this map in their average) and its standard error for each map per conditions and age groups.

